# Tracking changes in behavioural dynamics using prediction error

**DOI:** 10.1101/2020.10.19.346080

**Authors:** Tom Lorimer, Rachel Goodridge, Antonia K. Bock, Vitul Agarwal, Erik Saberski, George Sugihara, Scott A. Rifkin

## Abstract

Automated analysis of video can now generate extensive time series of pose and motion in freely-moving organisms. This requires new quantitative tools to characterize behavioural dynamics. For the model roundworm *Caenorhabditis elegans*, body pose can be accurately quantified from video as coordinates in a single low-dimensional space. We focus on this well-established case as an illustrative example and propose a method to reveal subtle variations in behaviour at high time resolution. Our data-driven method, based on empirical dynamic modeling, quantifies behavioural change as prediction error with respect to a time-delay-embedded ‘attractor’ of behavioural dynamics. Because this attractor is constructed from a user-specified reference data set, the approach can be tailored to specific behaviours of interest at the individual or group level. We validate the approach by detecting small changes in the movement dynamics of *C. elegans* at the initiation and completion of delta turns. We then examine an escape response initiated by an aversive stimulus and find that the method can track return to baseline behaviour in individual worms and reveal variations in the escape response between worms. We suggest that this general approach – defining dynamic behaviours using reference attractors and quantifying dynamic changes using prediction error – may be of broad interest and relevance to behavioural researchers working with video-derived time series.

## Introduction

Behaviour mediates individual interaction with the outside world. A systematic description of behaviour is therefore key to linking dynamic internal (e.g. neural) states, with external biotic and abiotic conditions in natural situations. However, even in cases where behaviour can be easily observed, finding a simple quantitative dynamic description is challenging. Increasingly, motion and pose of freely-moving organisms can be automatically tracked and quantified using computer vision (e.g. ***Mathis et al., 2018***). The model roundworm, *C. elegans*, presents a particularly well-established, promising and tractable case: its free motion on a surface can be automatically quantified from video in exceptional detail using only a few coordinates (***Stephens et al., 2008***; ***Brown et al., 2013***; ***Broekmans et al., 2016***), and its neuronal network is topologically stereotyped and contains only 302 neurons (***White et al., 1986***). Here we address the challenge of quantitatively defining and comparing behavioural dynamics in time series data, focusing on *C. elegans* locomotion as a case study.

On agar, *C. elegans* locomotion is characterized by forward motion with intermittent brief reversals. 35 years (***Chalfie et al., 1985***) of studies involving genetic perturbations, genetic sensors, electrophysiology, laser ablation, electron microscopy, computational modeling and other tools have mapped the connectome underlying this behaviour, clarified the contributions of allelic differences to behavioural variation, and have quantitatively dissected how locomotory decisions arise from signaling between network components (reviewed in ***Zhen and Samuel (2015)***). Here we focus at the organism level, on the locomotion itself, following a series of papers over the last 12 years that have developed a quantitative framework for describing and analyzing worm movement.

This quantitative framework for *C. elegans* movement on a surface relies on the observation that freely moving worm body poses can be represented sufficiently as coordinates in a 4 to 6 dimensional space of *eigenworms* (eigenvectors of worm body segment angles) (***Stephens et al., 2008***; ***Brown et al., 2013***; ***Broekmans et al., 2016***). This low dimensional representation can be linked back to traditional categorical descriptions such as direction of motion and turn type or can provide a finer-grained dictionary of pose motifs (e.g. ***Brown et al., 2013***). From such approaches, previously unclassified distinctions in behaviour and novel characteristics of mutants (e.g. ***Broekmans et al., 2016***; ***Brown et al., 2013***) demonstrate the power of this near-complete quantitative description of movement.

Categorizing worm behaviour by pose or motion (e.g. ***Brown et al., 2013***; ***Schwarz et al., 2015***; ***Szigeti et al., 2015***; ***Fukunaga and Iwasaki, 2017***; ***Javer et al., 2018***, ***2019***) can provide essential information regarding worm strain or internal state. This information can be exploited through the use of parametric or non-parametric models. For example, timing of reversals in direction (***Stephens et al., 2011***) and escape speed (***Daniels et al., 2019***) have been successfully predicted using parametric models and behavioural data alone (i.e. without using neural activity data). Recently, nonparametric models, which may make fewer assumptions, have shown great success whether including neural input (e.g. ***Brennan and Proekt, 2019***) or using body motion alone (e.g. ***Costa et al., 2019***; ***Ahamed et al., 2019***). ***Costa et al. (2019)*** used a hybrid approach in which eigenworm time series were dynamically partitioned and approximated locally with a sequence of linear models, revealing larger variation within behavioural types than previously thought. Recent papers have exploited delay-coordinate embeddings to reconstruct an approximate state space (in the dynamical systems sense) in which present states maximally determine future states. ***Ahamed et al. (2019)*** used this to obtain a comprehensive characterisation of the nonlinear dynamics of *C. elegans* locomotion. ***Brennan and Proekt (2019)***, by incorporating neural dynamics, were able to predict behavioural transitions up to 30 seconds in advance. A defining feature underlying the success of these nonparametric, embedding-based approaches is their ability to exploit the nonlinear dynamics evident in natural data of this kind, without requiring that a simple functional form can be found to accurately describe those dynamics (***Sugihara and May, 1990***).

Because even genetically identical individual worms can show distinct yet self-consistent patterns of motion (***Yemini et al., 2013***; ***Brennan and Proekt, 2019***), it can be difficult to measure subtle changes in behaviour that may reflect important changes in internal state. Categorical definitions of movement or pose behaviour (e.g. ***Schwarz et al., 2015***), however fine-grained (e.g. ***Brown et al., 2013***), necessarily collapse subtle variation onto a dictionary of stereotypes. Non-categorical distinctions of behaviour must necessarily confront the problem of measuring distances between time series – a highly non-trivial problem. The recent success of dynamic embedding representations of worm motion (***Ahamed et al., 2019***; ***Brennan and Proekt, 2019***), suggests that this problem could be addressed within the framework of empirical dynamic modeling (EDM) (***Sugihara and May, 1990***).

EDM relies on reconstructing attractors that encode system dynamics directly from time series data (for a simple introduction, see [Introduction to Empirical Dynamic Modeling](https://www.youtube.com/watch?v=fevurdpiRYg)). In the context of worm motion, attractors can be reconstructed from data agnostically, i.e. describing the dynamics in a given set of data without requiring prior assumptions about worm, strain, or motion type. These attractors thereby provide a definition of worm dynamics for a given set of observational data to which other worm dynamics can be easily compared.

In this manuscript, we assess the feasibility and robustness of extending eigenworm representations of motion to dynamic attractors based on time-delay-coordinate embeddings. We then exploit this observed robustness to propose a prediction-based scheme for detecting dynamic change. We validate our scheme using the fluctuations of motion dynamics inherent to *delta turns* (***Broekmans et al., 2016***) and show that our scheme provides a novel, informative and nuanced view of the return to baseline worm motion following an aversive stimulus.

### Method overview

To quantitatively assess changes in worm motion dynamics, we require a quantitative representation of those dynamics. We construct this representation based on a time-delay embedding of a reference time series, or library, of observed worm poses (Fig. 1). Each point in this embedding corresponds to a short sequence, of length *E*, of successive worm body poses in the library, at regular time intervals *τ* (Fig. 1A). Note that because we encode each worm pose here using five eigenworm coefficients, the embedding space has 5*E* dimensions in total. The dynamic information about the order in which these pose sequences occur within the library defines a trajectory through the embedding space (Fig. 1B). With an appropriate choice of *E* and *τ*, the trajectory unfolds into a nearly-repeating but non-intersecting structure that we here call an attractor.

**Figure 1.**
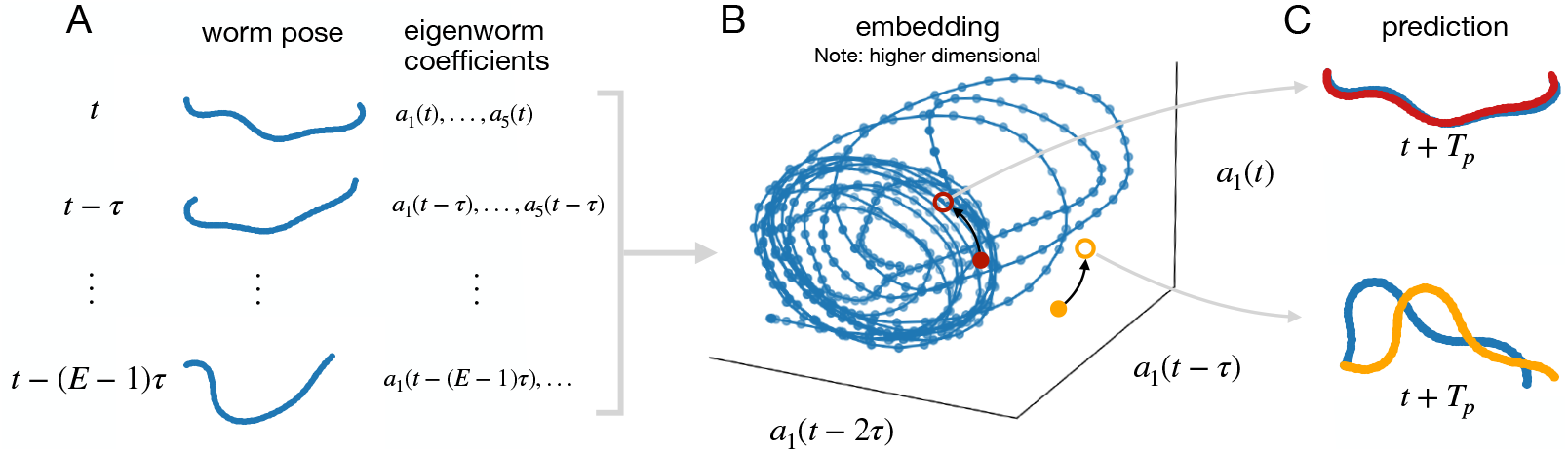
Embedding and predicting worm pose time series to quantify dynamic similarity (schematic). A) Static worm poses (encoded as 5 eigenworm coefficients) from a reference library of worm behaviour are embedded in their dynamic context: short sub-sequences of *E* worm poses. B) These embedded points, together with their temporal sequence, define a trajectory. An appropriately-embedded time series resolves into an attractor. By following nearby points on the attractor, predictions can be made for out-of-sample data. C) Out-of-sample data from dynamics similar to the reference library result in good predictions (red), whereas different dynamic regimes will be predicted poorly (orange).

The dynamic information contained in such an attractor can be exploited to make predictions. Nearby points (where each point is a sequence of *E* worm poses) on the attractor belong to locally similar trajectories (Fig. 1B). This means the trajectory of an out-of-sample point can be estimated by following nearby points on the library attractor as they move forward in time (see ***Sugihara and May, 1990***; ***Sugihara, 1994***). An out-of-sample point originating from dynamics that are similar to those encoded in the library attractor should permit a better prediction than an out-of-sample point originating from different dynamics (Fig. 1C). Thus, out-of-sample prediction accuracy may function as a useful measure of dynamic similarity (cf. ***Liu et al., 2012***).

We wish to maximise the temporal resolution of our method, in order to precisely localise behavioural changes in time. Therefore we set the embedding-time-delay interval *τ* and the prediction horizon *T_p_* equal to the sampling interval of the data. For the forward prediction, we use the S-Map method (***Sugihara, 1994***) as it allows the sensitivity to local attractor structure to be tuned, thereby optimally exploiting nonlinearity in the data. This leaves us with two free parameters that define our measure: the number of successive worm poses *E*, and the S-Map nonlinearity parameter *θ* (see also Methods and Materials).

## Results

### Parameter robustness

A natural concern is whether such a measure will be robust enough to parameter variation to actually be useful. We assess this on a suite of publicly available worm motion data from four strains: wild type (N2) and three diverse mutants (*octr-1(ok317)*, *unc-80(e1069)* and *dpy-20(e1282)IV*), using library and prediction intervals of 1000 samples each (Fig. 2, see Methods and Materials).

**Figure 2.**
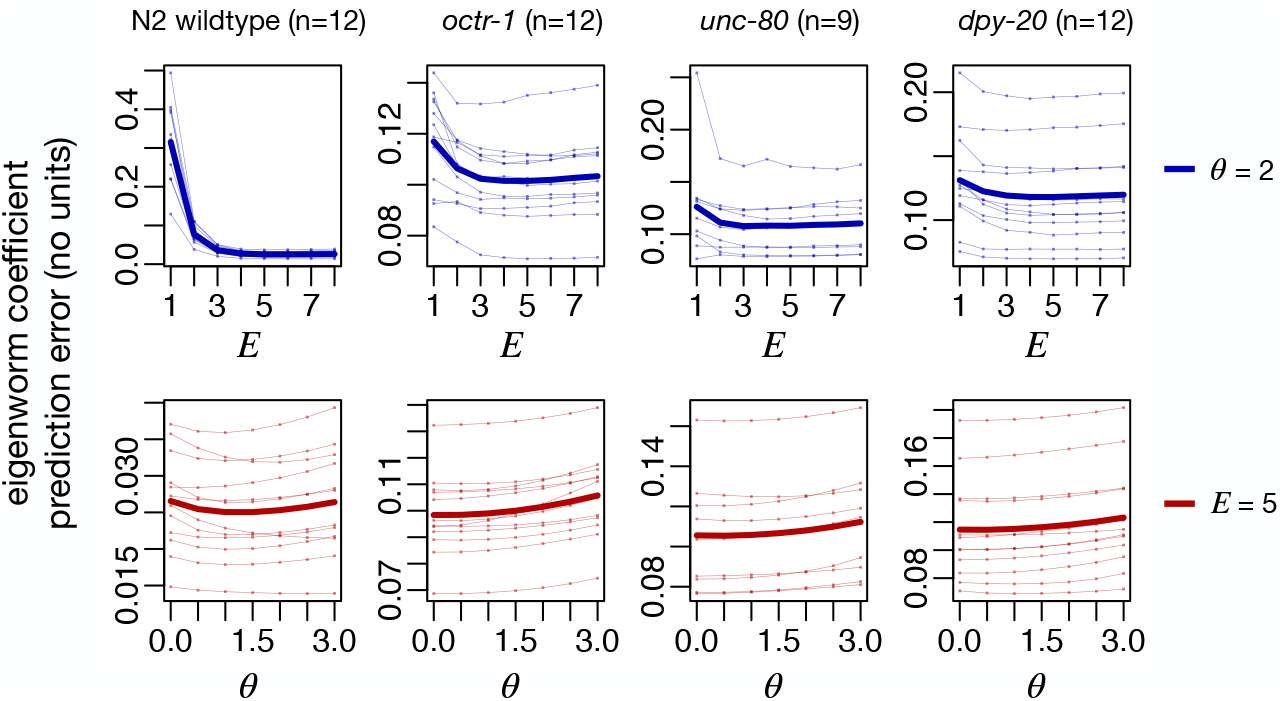
Prediction error is robust to parameter variation. Top row: robustness to *E* (at *θ* = 2) for data from a wild type and three mutants (headings; see text). Bottom row: robustness to *θ* (at *E* = 5) for the same four worm types as top row. Thin lines are individual worms, thick lines are means. Note that there is a coordinate system variation between the wildtype and mutant worm data, which hinders direct comparison of error magnitudes.

Average prediction error remains stable as a function of both *E* and *θ*, over a wide range of parameter settings, for almost all worms of each type (Fig. 2). Within each type of worm, the shape of the error as a function of the parameters shows strong consistency between individuals. Generally, the optimal parameter settings occur at *E* > 1, and the wildtype shows evidence of nonlinearity, i.e., the optimal value of *θ* > 0, which is consistent with previous observations (***Ahamed et al., 2019***; ***Costa et al., 2019***). The mutant worms on the other hand generally do not show clear evidence of nonlinearity on average, though the sensitivity to *θ* is very weak over the parameter range explored. This more linear characteristic of the mutant data may be attributed to at least two causes: 1) the mutant data do not contain self-intersecting worm poses, whereas these were reconstructed in a second processing step for the N2 wildtype data ***Broekmans et al. (2016)***; 2) the sampling rate of the mutant worm data is almost twice that of the N2 wildtype data (30 Hz vs. 16 Hz), which will make the data appear locally far more linear at the single timestep scale. Indeed, looking at an additional wildtype from the same data source as the mutant time series (i.e., sampled at 30 Hz without self-intersecting turns) reveals a very similar *E*, *θ* robustness profile as the other mutants (Fig. 2–Fig. supp. 1). Despite all of these differences between the acquisition and pre-processing of the time series however, the prediction error remains stable over a single wide parameter range across all of these data. Furthermore, the prediction error is robust in detail through time, with the error time series showing strong correlation across parameter settings (Fig. 2–Fig. supp. 2). For all subsequent analyses, we therefore choose mid-range parameters *E* = 5 and *θ* = 2, to allow for any transient highly nonlinear behaviour that may occur, without substantially sacrificing average prediction error. In all subsequent analyses also, we work only with the N2 wildtype time series from ***Broekmans et al. (2016)***. We therefore convert the observed and predicted eigenworm coefficients to worm body section angles (***Stephens et al., 2008***) prior to calculating error, to allow the use of intuitive error units: RMS error of body section angles in radians.

### Validation and dynamic sensitivity

In addition to being stable, we also require our measure to be meaningful: to detect subtle variations in worm motion dynamics that may potentially have an exogenous origin, such as a change in (latent) internal state or external conditions. We propose here that delta turns (***Broekmans et al., 2016***) may provide an ideally constrained example of such exogenous interference. During delta turns, where the worm must intersect and then cross over (or under) its own body, the initiation and completion of the turn may result in both worm body shape deformation (thereby temporarily modifying the coordinate system of the dynamics) and stick-slip-type dynamics that are likely to be highly variable. Furthermore, because the traditional algorithms used to extract worm pose from image data fail when the worm body intersects itself, a supplementary inference method is needed (***Broekmans et al., 2016***). The uncertainty inherent in making such an inference results in a small increase in worm pose estimation error (***Broekmans et al., 2016***). This error is exogenous to the natural dynamics of worm motion. Taken together, we thus we expect that a method for detecting variation in worm dynamics should show a distinct spike during the transition from non-intersecting pose to crossing pose (at delta turn initiation), and from crossing to non-intersecting (at delta turn completion).

In Figure 3, the time series of a foraging worm from the ***Broekmans et al. (2016)*** data (see Methods and Materials) do not show obvious differences between the reference library and prediction intervals (Fig. 3A). Accordingly the prediction based on our attractor performs well at almost all time points, with the exception of four pronounced pairs of sharp peaks and one isolated sharp peak (Fig. 3B). The location of these peaks does not correspond to any clear signal change in the original time series (arrows, Fig. 3A). The time points where the magnitude of the third eigenworm coeffcient |*a*_3_| > 15 correspond to four tight turns (***Broekmans et al., 2016***) that upon visual inspection are confirmed to be delta turns (red points, Fig. 3B). The four delta turns are each book-ended by pronounced spikes in error, that correspond to the transitions during initiation and completion of the delta turn described above (Fig. 3C). Closer inspection of the localisation of prediction error along the worm body confirms that the head-end of the worm is responsible for the error peak during delta turn initiation, and the tail-end during delta turn completion in all four cases, exactly as would be expected (Fig. 4). The origin of the isolated spike in error around 8 seconds is thought to arise from a dynamic sequence during a ‘w’-shaped reverse motion that is sufficiently different from motions sampled in the library (see also Discussion).

**Figure 3.**
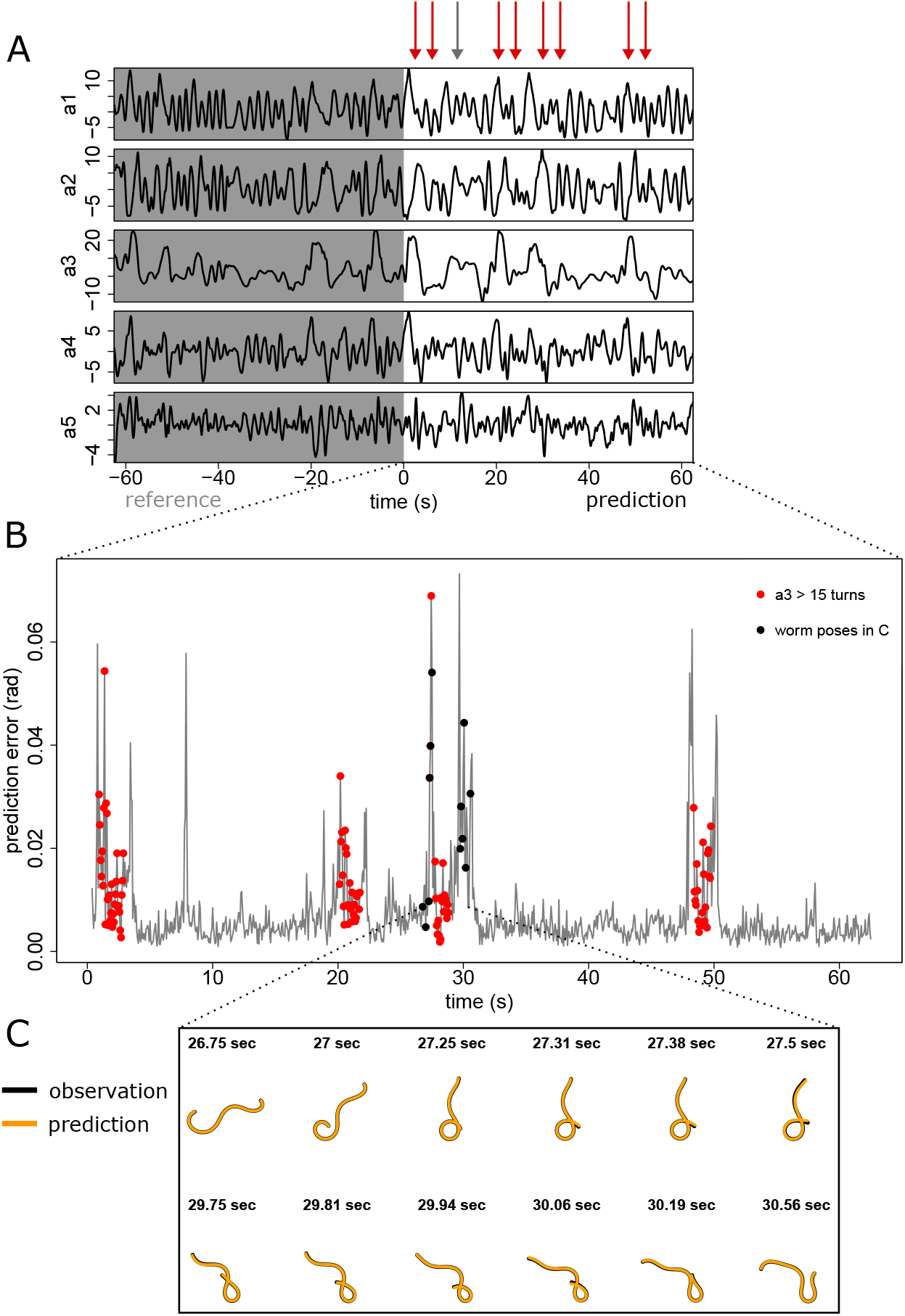
Prediction error reveals anomalous dynamics in worm pose time series. A) Original time series of eigenworm coefficients indicating the split into reference library and prediction set. Error peaks in (B) indicated by arrows do not correspond to obvious features in the time series. B) Prediction error corresponding to prediction set in (A). Turns (red dots), characterised here by third eigenworm coeffcient 15, are book-ended by periods of anomalous dynamics. The mean error for a constant predictor (i.e.|*a*_3_| > the prediction that the worm pose is the same as the previous time step) is higher than the vertical axis range. C) A closer look at the worm pose time series (black) and predictions (orange) reveals that the anomalous dynamics correspond to delta turn initiation and completion, when the worm head and tail respectively transition between self-intersection and no self-intersection.

**Figure 4.**
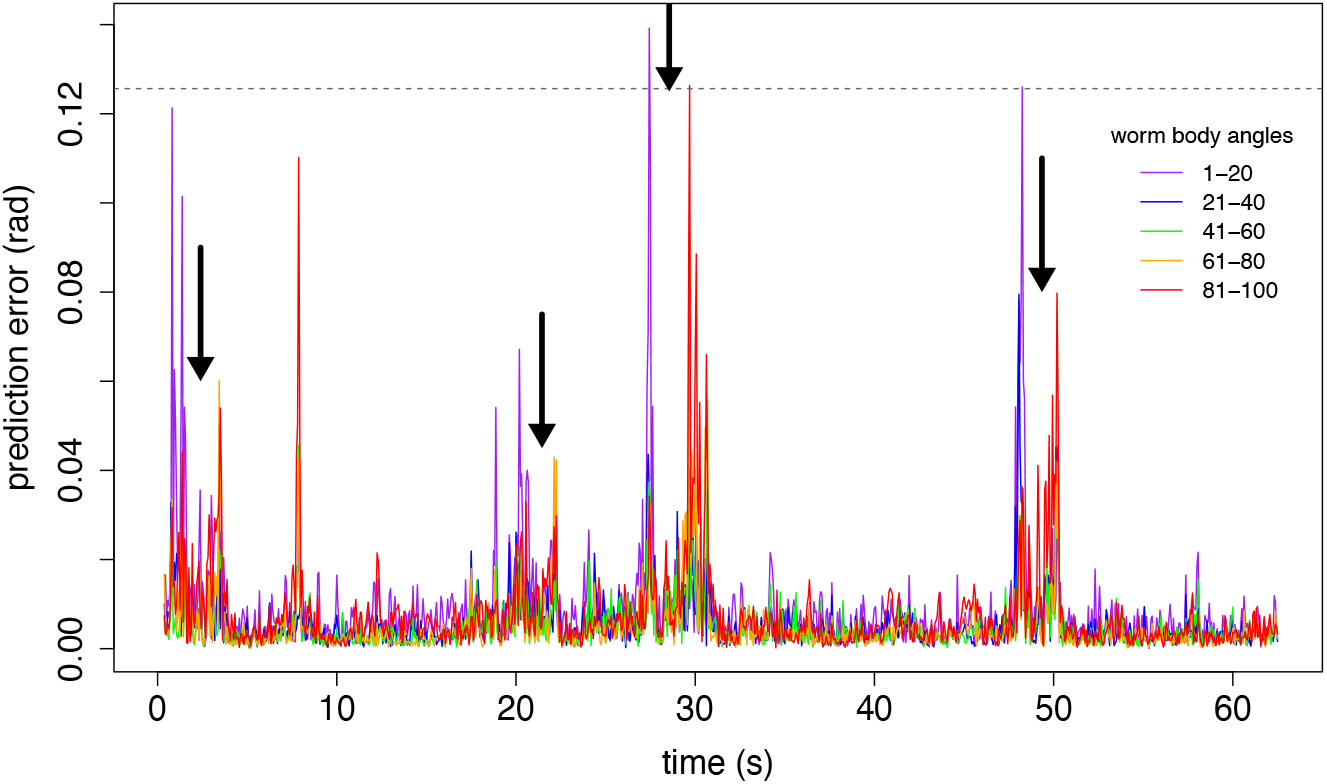
Systematic localisation of prediction error along the worm body is consistent with delta turn self-interference. The four major delta turns around 2, 20, 30 and 50 seconds all show the same characteristic pattern of head-localised error at turn initiation (angles 1–20), and tail-localised error at turn completion (angles 81–100). During the middle portion of the turn, the dynamics are more predictable. Dashed horizontal line indicates the mean whole-worm error for the constant predictor (i.e. predict that the worm pose is the same as the previous time step).

### Application example

We use the measure of worm dynamics variation to examine the return to baseline behaviour following an aversive stimulus. In this example, described in ***Stephens et al. (2008)***, worms were subjected to a brief infra-red laser pulse to the head after 10 seconds of recording. The worms then exhibited a characteristic escape response: rapid reverse crawling motion, followed by a tight (omega) turn, and then forward motion in the new direction away from the location of the laser pulse (***Stephens et al., 2008***).

We use the first 80% of time points before the stimulus as a reference library of baseline behaviour, to allow us to compare out-of-sample prediction error before and after the stimulus. We do not perform any additional optimisation of the parameters of our algorithm, taking instead the same parameters used earlier: *E* = 5 and *θ* = 2 (see Methods and Materials). We do this despite the fact that these escape response datasets have a different sampling rate from the foraging worm datasets (20 Hz vs. 16 Hz) in order to highlight the practical robustness of our method. Because the escape response contains behaviours that are almost surely not contained in the pre-shock reference library of worm dynamics, we perform an approximate classification of worm behaviour across the entire time series to aid interpretation. We identify forwards and backward crawling motion based on the sign of the apparent phase velocity in the (*a*_1_*, a*_2_) plane (***Stephens et al., 2008***) (provided the amplitude in this plane is sufficiently large, see Methods and Materials), and we conservatively identify as putative tight turns time points where |*a*_3_| > 10.

Figure 5 shows the typical escape-response sequence of reverse–turn–forward motion following the stimulus. This sequence is also reflected in the prediction error time series: at stimulus onset, there is a sharp increase in the prediction error that persists throughout the rapid reverse crawling and subsequent turn, and then eventually returns close to baseline during subsequent forward motion (e.g. Fig. 5A,B). In some cases (e.g. Fig. 5D,E), the prediction error remains high, suggesting that the movement dynamics of the worm towards the end of the time series remains substantially different from the movement dynamics prior to the shock. However, care is required when interpreting this result. By definition, baseline is whatever dynamics occupy the reference library. If the reference library does not adequately sample the behavioural regime of interest, then it can not adequately represent it. For these escape response data, there are very few time points available before the shock from which to construct a reference library, and so the reference library is short: 160 time points, or 8 seconds (by contrast, for the foraging worms we used 1000 time points, or 62.5 seconds). Nonetheless, even using a limited library, in Fig. 5A,B,F,G and H, the prediction error clearly indicates that the forward crawling motions at the beginning and the end of the sequence are similar (despite differences in phase velocity) and that they are substantially different from the bulk of the escape response.

**Figure 5.**
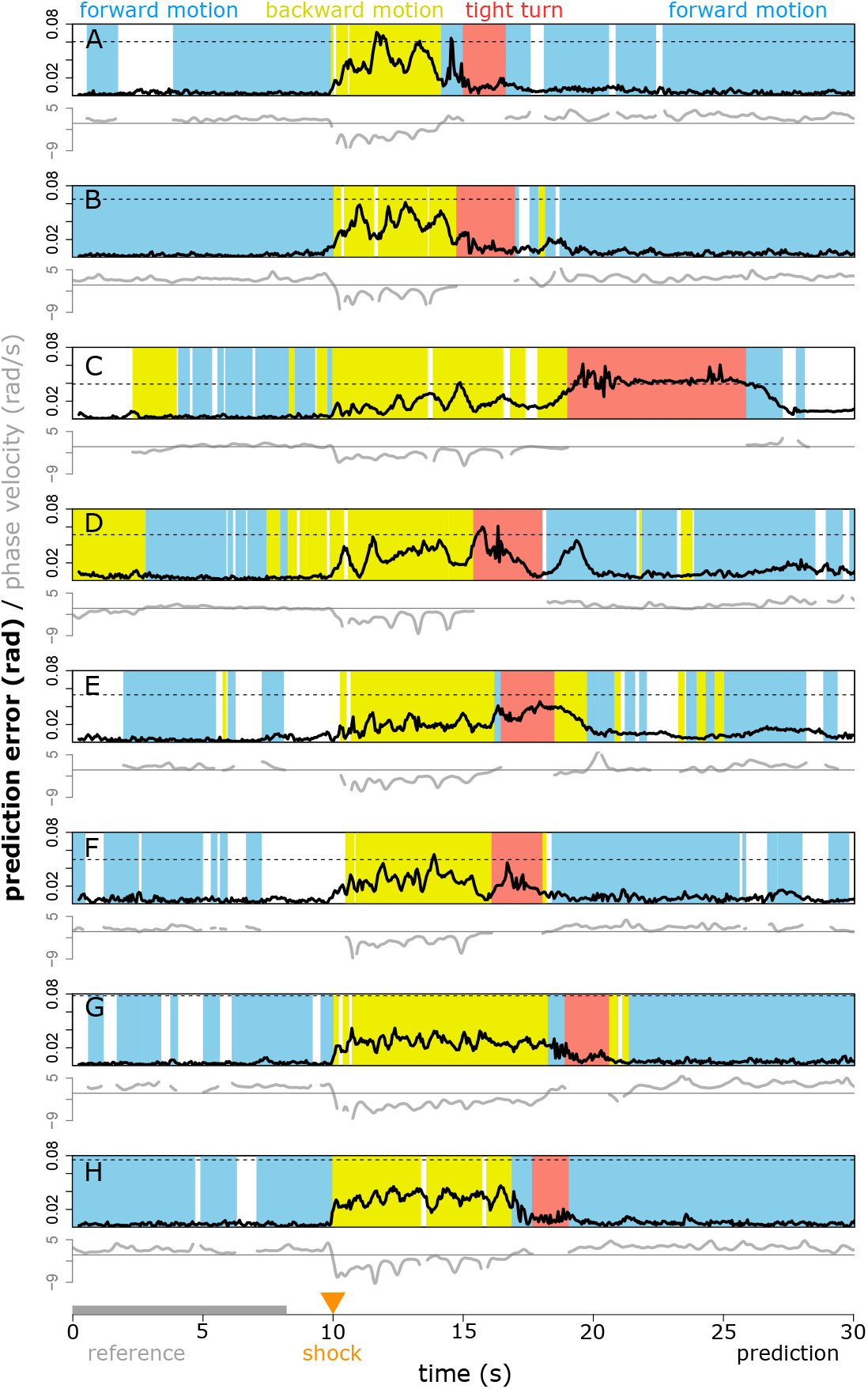
Prediction error (black lines) can successfully identify the departure from and return to baseline behaviour following an aversive stimulus at 10s, despite a very small library size (first 8 seconds, grey). Different rows correspond to different worms (see Materials and Methods). Background colours indicate approximate behavioural classification into tight turns (red, |*a*_3_| > 10), forward and backward motion (blue and yellow, positive and negative phase velocity in the *a*_1_*, a*_2_ plane, respectively), and unclassified (white, when amplitude in the *a*_1_*, a*_2_ plane or the estimated phase velocity are too small), intended as an approximate guide only (see Appendix 1). Grey lines indicate the apparent relative phase velocity in the *a*_1_*, a*_2_ plane used in the classification scheme (see Appendix 1). The dashed horizontal lines indicate the mean error for the constant predictor (i.e. predict that the worm pose is the same as at the previous time step) provided that error lies within the vertical axis range.

Comparing between escaping worms reveals fine-grained variations in escape response. If we use, for example, the worm in Fig. 5A as the reference library for the worm in Fig. 5B, and vice versa (Fig. 6), we see a dramatic difference in the escape response of the worm in Fig. 5A around 15 seconds (Fig. 6B). This behavioural sequence, that is different from any sequence exhibited by the reference worm, results in a peak error several times higher than other major peaks in either error time series. This anomalous behaviour is also visible in the approximate classification scheme: the apparent (*a*_1_*, a*_2_) plane phase velocity changes sign between the escape reversal and the turn, resulting in a transient ‘forward motion’ classification. Importantly, however, detecting this difference using our method does not require any special foresight about which data features (e.g. phase velocity) should be used for detection. In addition to this most obvious feature, we see clearly that the forward motions of each worm before and after the escape response sequence are broadly similar (low error, Fig. 6A,B). The error during the initial reversal shows similar behaviour in each case, with an initial peak of similar magnitude, that then declines throughout the reversal sequence. This highlights a challenge in the measurement of worm motion behaviour: in most dynamic representations (e.g. ***Ahamed et al., 2019***; ***Brown et al., 2013***) rate and shape are coupled. That is, if a worm were to run through exactly the same sequence of poses at a different rate, this would map into a different position in the dynamic representation (embedding) space. This is also true of our method. Hence, during the rapid reversal, which is initially fast and gradually decays in speed and phase velocity (Fig. 5 and ***Daniels et al., 2019***), we see a concurrent decrease in prediction error as motion slows (except at the end of the reversal in Fig. 6A, where the reference worm’s behaviour differed). Note however, that the prediction error during reversal when predicting from one worm to the other is still substantially lower than the prediction error during reversal in Fig. 5, where no reversal time sequence was included in the library.

**Figure 6.**
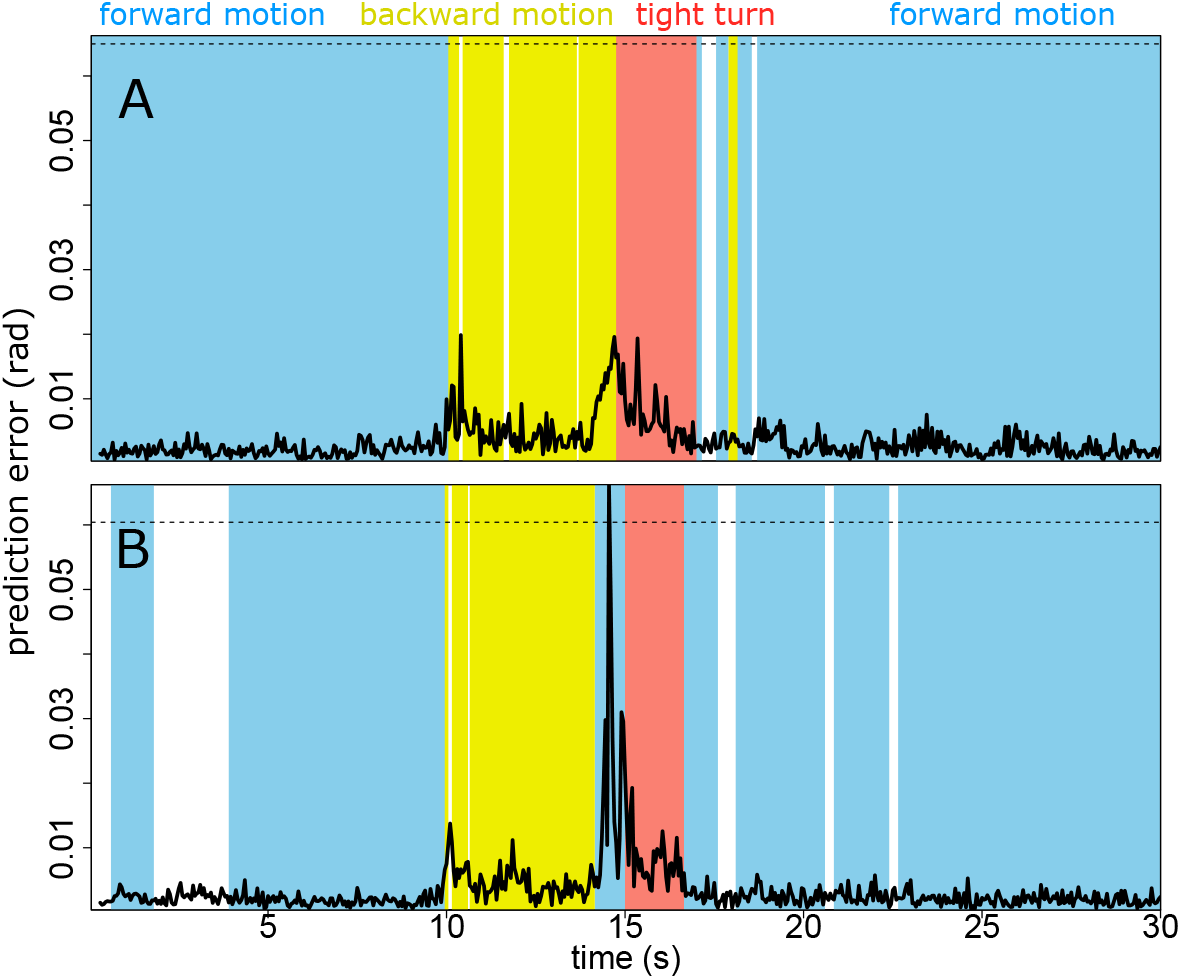
Predicting from one escaping worm to another reveals differences in escape response. Using the escaping worm in Fig. 5A as the reference library for the escaping worm in Fig. 5B shows that behavioural dynamics exhibited by the predicted worm are broadly similar to examples from the reference worm (A). Predicting the other way, however (B), reveals that the worm in Fig. 5A displays a behaviour around 15 seconds that is inconsistent with the reference library from the worm in Fig. 5B. This is further confirmed by a change phase velocity sign (apparent ‘forward motion’). Other parts of the escape response, as well as the forward motion before and afterward are similar. Dashed horizontal line indicates the mean error for constant the predictor (i.e. predict that the worm pose is the same as at the previous time step).

## Discussion

The approach presented here exhibits several advantages in comparison with other methods that can be used to detect change in worm motion dynamics. Figures 3, 4, and 5 show that the output signal of the algorithm lets us localize changes in worm behaviour to within a few time samples. This property is a direct consequence of our use of prediction. Template-matching approaches to measure motion change (e.g. ***Brown et al., 2013***) are similar on a local scale to time-delay embeddings. However, because these approaches do not use the dynamic trajectory information in an embedded space, they do not fully exploit the time resolution of the data. Comparing predictive output at a single time point focuses this information to single-sample resolution. In order for such fine scale prediction to be meaningful, however, it has to be able to exploit relevant information globally in time—achieved here using the attractor. This data-driven focusing of relevant information across time scales is a natural advantage of the EDM approach.

A second advantage of our approach is its flexibility and sensitivity. These arise from the use of a user-specified library for the construction of the attractor, allowing the relevant dynamic reference information to be tailored with great precision. Although here we have focused on dynamic change within and between individual worms, the approach can be easily and naturally extended to composite libraries encompassing broader classes of behaviour.

This flexibility and sensitivity comes with a cost. An inappropriate choice of reference library can lead to output that carries little useful information (see Fig. 5D,E). We suggest, therefore, the use of an independent method of behavioural categorisation (such as the one used here) in conjunction with our method, to aid the user in distinguishing dynamic differences due to behaviours that were absent or under-sampled in the reference library from dynamic differences due to more subtle changes in state (see Appendix 2). In some applications, however, it may not be necessary to make this distinction. For example, methods that look at average behavioural differences over an extended time window to give a global measure of behavioural difference will usually conflate these two types of difference but may still yield useful information (e.g. ***Brown et al., 2013***).

Because our method uses a fixed sampling interval to define the embedding, pose sequences performed at different rates will be represented at different locations in the embedding space. Although the method appears robust (see Fig. 2 and the adoption of parameters from Fig. 3 in Figs. 5 and 6 despite a change in sampling rate) it may be desirable in some instances to disentangle the rate of progression through a pose sequence from the pose sequence itself. This might allow, for example, the differences in worm escape response reversal to be determined more precisely (see Fig. 6). To achieve such a separation would require the time delay embedding construction to be adjusted according to, e.g. phase velocity or worm speed: a process that would likely benefit from an additional dimensionality reduction step in time series embedding such as that used by ***Ahamed et al. (2019)***. Here, we have focused on the simplest embedding scheme to aid interpretability.

In cases where the sampling rate of the data is very high (e.g. 30 Hz and above), it can be expected that the constant predictor will become quite accurate, as the pose change between successive frames becomes very small. (Note the substantially lower constant predictor error in Figs. 5 and 6 where the sampling rate is 20 Hz, than in Figs. 3 and 4 where the sampling rate is 16 Hz.) In some such cases, it may be desirable to sacrifice temporal resolution for increased sensitivity, by setting *τ* and *T_p_* to twice the sampling interval of the data. In this initial study we have focused on maximum temporal resolution as a base case.

More generally, the simple, flexible, sensitive and robust characteristics of our behavioural change measure when analyzing *C. elegans* pose dynamics result from the ability of EDM tools to exploit the exceptional richness and resolution of time series data, without imposing strong model assumptions. The recent success of other empirical nonlinear dynamics approaches (***Brennan and Proekt, 2019***; ***Ahamed et al., 2019***) suggests that such techniques may prove increasingly useful in future *C. elegans* behavioural analyses.

Although we have focused on *C. elegans* here, the embedding and prediction tools that underlie our approach are applied widely to time series data in general. The key observation that prediction error can be used to measure meaningful behavioural change can therefore also be explored in the increasing number of other high-resolution behavioural time series extracted from video (e.g. ***Mathis et al., 2018***).

## Methods and Materials

### Data

We use the previously-published foraging and escape time series of ***Broekmans et al. (2016)***. The foraging dataset consists of time series of the first five eigenworms from twelve N2 wildtype worms. Continuous uninterrupted blocks of 2000 samples (16Hz) were identified to avoid NaN values in the original data as follows: worm 1: 10001 to 12000, worm 2: 10001 to 12000, worm 3: 8501 to 10500, worm 4: 10501 to 12500, worm 5: 10501 to 12500, worm 6: 10501 to 12500, worm 7: 10501 to 12500, worm 8: 10501 to 12500, worm 9: 11001 to 13000, worm 10: 11001 to 13000, worm 11: 11001 to 13000, and worm 12: 11001 to 13000. For the robustness calculations (Fig. 2), these intervals are extended slightly due to the embedding construction method used (see Embedding construction, below). The escape response dataset includes the full 5 eigenworm time series from ***Broekmans et al. (2016)***, measured for 30 seconds at 20 Hz. Time series in Fig. 5 A to H correspond to escaping worm numbers 50, 69, 37, 76, 2, 5, 8 and 12 respectively in the original dataset.

In our robustness analysis we include time series of the first five eigenworms from three additional mutant strain datasets from [OpenWorm](http://movement.openworm.org/) (***Szigeti et al., 2014***), collected as per ***Yemini et al. (2013)***. The mutations selected are *octr-1(ok317)* (strain VC224), *unc-80(e1069)V* (strain CB1069), and *dpy-20(e1282)IV* (strain CB1282). Twelve individual worms were analyzed for *octr-1(ok317)* and *dpy-20(e1282)IV*, while nine individuals were analyzed for *unc-80(e1069)V*. Embeddings were constructed using the discontinuous method (see below), starting from sample 1001, and generating embeddings of 1000 points each for the library and prediction. See Appendix 1 for lists of the specific files included.

### Analysis

The code used in our analysis is available via GitHub [here](https://rachelgoodridge.github.io/c_elegans_pred_error/).

### Embedding construction

We use two slightly different methods to construct embeddings: a continuous method and a discontinuous method.

The *continuous method* is suitable for time series data that do not contain substantial breaks (i.e. NaN values). It generates embeddings from fixed data intervals (e.g. from sample 12001 to 13000 inclusive). This allows precise delineation of behaviours of interest, and simple identification of prediction error with time. Note that because this method only has access to exactly the data interval specified, the embedding coordinates for the first few time points will be undefined, as time lags from before the interval are not available. The continuous method is used for all figures where error time-series are plotted for the N2 wildtype worms: Fig. 2–Fig. supp. 2; Figs. 3 to 6; Appendix 2.

The *discontinuous method* can account for breaks (NaN values) in the time series. It is used here for all figures that include the mutant worm data (which contain NaN values during self-intersecting worm poses): Fig. 2 and Fig. 2–Fig. supp. 1. The discontinuous method takes as input only a starting sample (e.g. sample 1001), and proceeds to construct an embedding from there, accepting only embedding points for which all lag coordinates and target coordinates are available. To avoid embedding mismatches due to changing *E* during robustness testing, we construct a master embedding for the highest value of *E*, and use subsets of this embedding for lower values of *E*.

#### Prediction and error metric

We use the [rEDM](https://github.com/SugiharaLab/rEDM) package in R. Using the S-map (***Sugihara, 1994***) feature in rEDM for each eigenworm coefficient and the *E* * 5 dimensional embedding, we make out-of-sample predictions of the first 5 eigenworm coefficients. In all cases we used *τ* = *T_p_* = M, where M is the sampling interval of the data. For quantification of prediction error for any given worm at a single time point, we convert predicted and observed eigenworm coefficients to 100 worm body section angles (***Stephens et al., 2008***), then calculate RMS error in radians, unless otherwise specified (cf. Robustness calculations, below).

#### Robustness calculations (Fig. 2)

We use the first 1000 embedded points (see Data and Embedding construction, above) to as the reference library attractor and predict from the remaining 1000 embedded points. The vertical axis of Fig. 2 shows the mean over time of the single time point RMS error between the observed and predicted eigenworm coefficients, for each worm. The thick lines described as “means” are simply constructed by taking the arithmetic mean of the values in the thin (individual worm) lines.

#### Time dependence of error (Figs. 3 to 6)

Figure 3 uses S-map analysis as above to make predictions on the eigenworm coefficients of foraging worm 1 from ***Broekmans et al. (2016)***, with parameters *E* = 5, *θ* = 2. The reference library is constructed from frames 10001–11000, while the eigenworm coefficient predictions are made using frames 11001–12000. Predicted and observed worm poses are displayed at points of interest by converting the body angles calculated using the eigenworms to coordinates in the plane assuming fixed length of each worm segment, and aligning predicted and observed worms at the head.

Figure 4 incorporates the same data and method as Figure 3, but with RMS error in predictions calculated over each of five equally-sized non-overlapping regions of the worm as indicated in the figure. These equally-sized regions are each composed of twenty body angles extracted from the eigenworm coefficients.

In Figure 5, we use the first eight seconds of pre-stimulus behaviour to predict post-stimulus escape response while tracking prediction error. For approximate behavioural classification, see Appendix 1.

In Figure 6, we proceed as per Figure 5, but instead of using the first 8 seconds of the predicted worm time series as the library, we use the entire time series of a different escaping worm, as specified in the text.

## Acknowledgments

This work was supported by: the Scripps Institution of Oceanography Postdoctoral Fellowhsip (TL); DoD-Strategic Environmental Research and Development Program 15 RC-2509, NSF DEB-1655203, NSF ABI-1667584, the McQuown Fund and the McQuown Chair in Natural Sciences, University of California, San Diego (GS); and NSF IOS-1936674 (SAR).

## Competing Interests

The authors declare no competing interests.

## Author Contributions

All co-authors contributed to all phases of the project. TL organized and led the final writing and analysis.

## Appendix 1

## Classification of worm behaviours

Given time series *a*_1_(*t*), *a*_2_(*t*), *a*_3_(*t*) corresponding to coefficients of the first, second and third eigenworms respectively, sampled at regular time intervals Δ*t* . Let *c*(*t*) be the class label assigned to the worm behaviour at time . We define our approximate classification scheme as follows (cf. ***Stephens et al., 2008; Broekmans et al., 2016***). Note that in the main text we have reversed the sign of the phase velocity relative to what is described here, such that forward motion corresponds to positive phase velocity.

**if** |*a*_3_(*t*) > 10 **then** *c*(*t*) =“tight turn”

**else if** 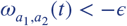 **and** 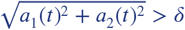 **then** *c*(*t*) = “forward motion”

**else if** 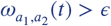 **and** 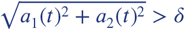 **then** *c*(*t*) = “backward motion”

**else** *c*(*t*) = “unclassified”

where 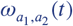 is a local estimate of phase velocity in the (*a*_1_*, a*_2_) plane given by:

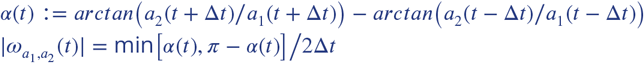

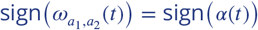 if *α*(*t*) < π − *α*(*t*), and

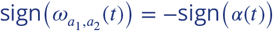 otherwise.

The minimum phase velocity threshold *ϵ* = 0.1 rad/s, and minimum phase plane amplitude *δ* = 3 (no units) were chosen to improve stability.

## Mutant worm data used in Figure 2

See separate Excel spreadsheet file *suppTableMutantWorms.xlsx*

## Appendix 2

## Classified library and prediction error time series for 12 foraging worms

**Appendix 2 Figure 1.**
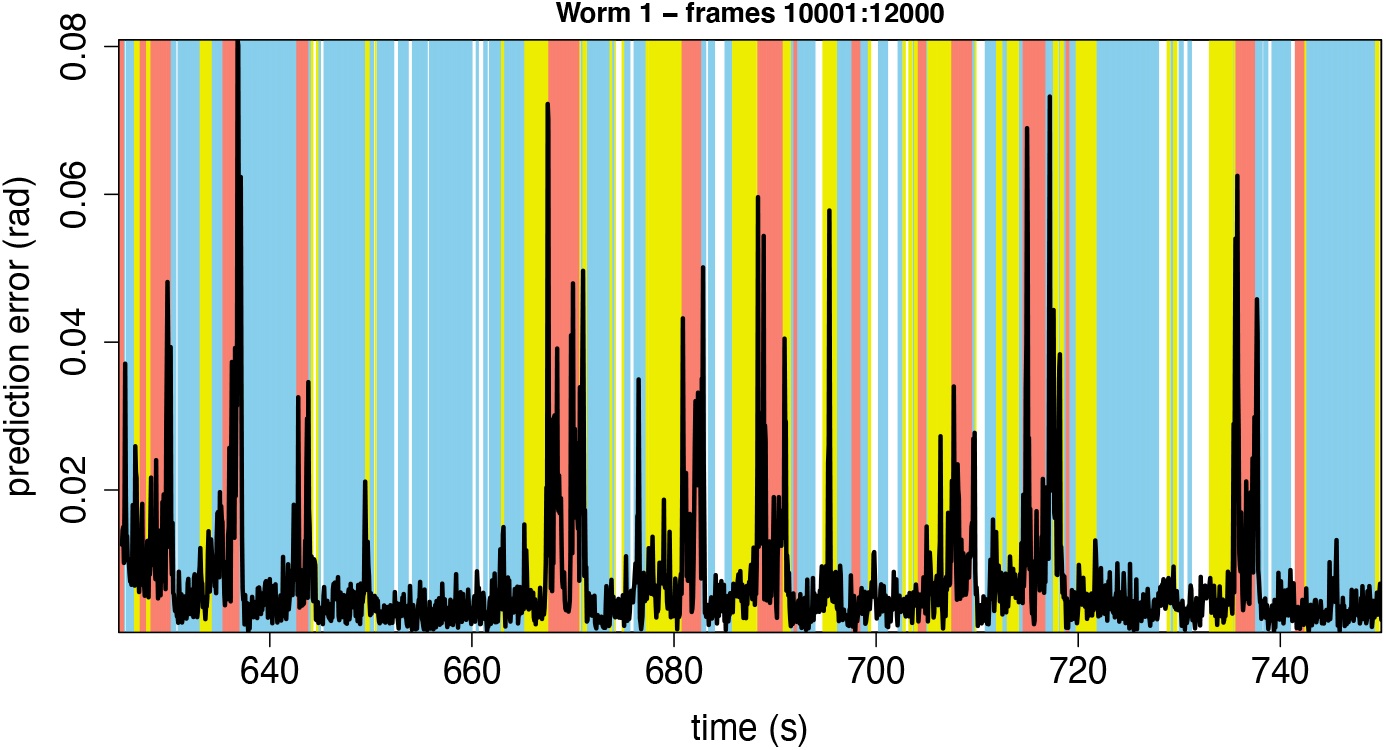
Full error time series for foraging worm 1 library (first half) and prediction (second half) intervals (see Fig. 3A). Background: approximate behavioural classification (blue, forward motion; yellow, backward motion; red, tight turn).

**Appendix 2 Figure 2.**
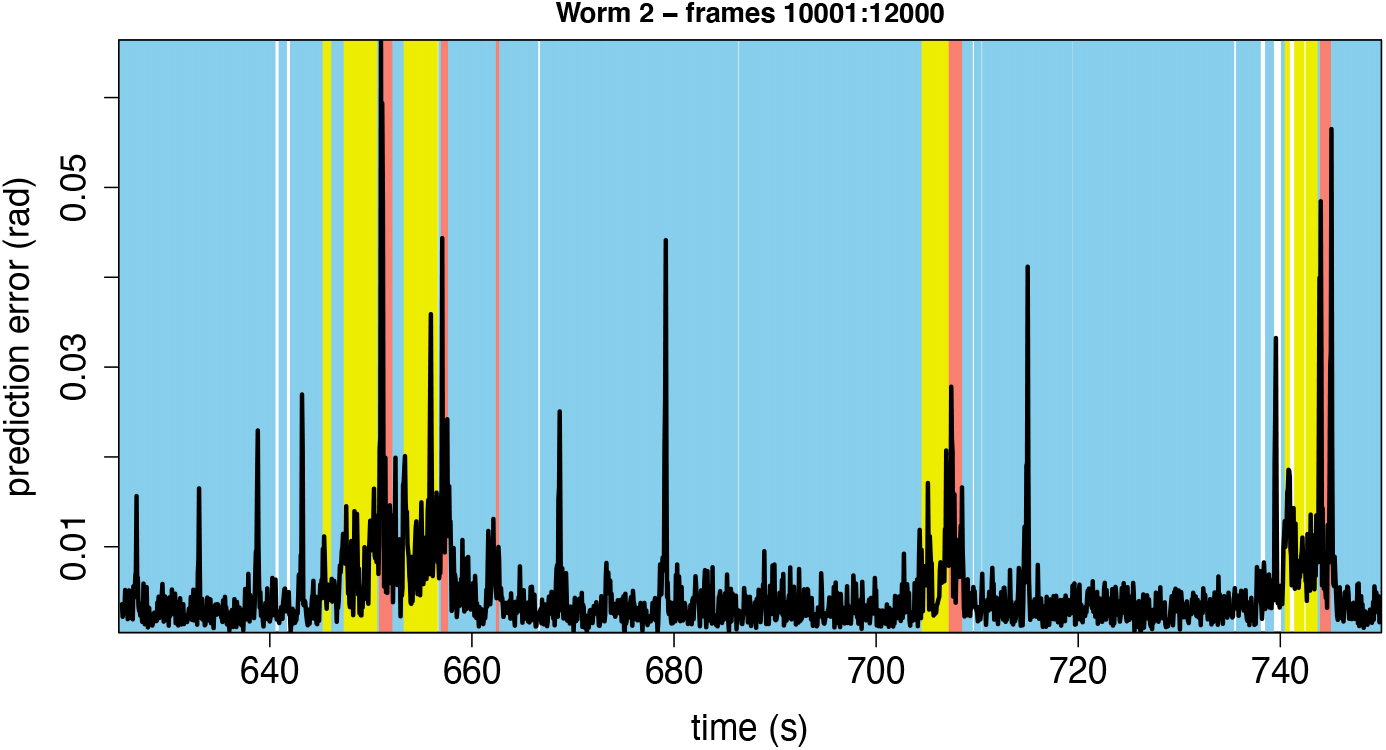
Full error time series for foraging worm 2 library (first half) and prediction (second half) intervals. Background: approximate behavioural classification (blue, forward motion; yellow, backward motion; red, tight turn).

**Appendix 2 Figure 3.**
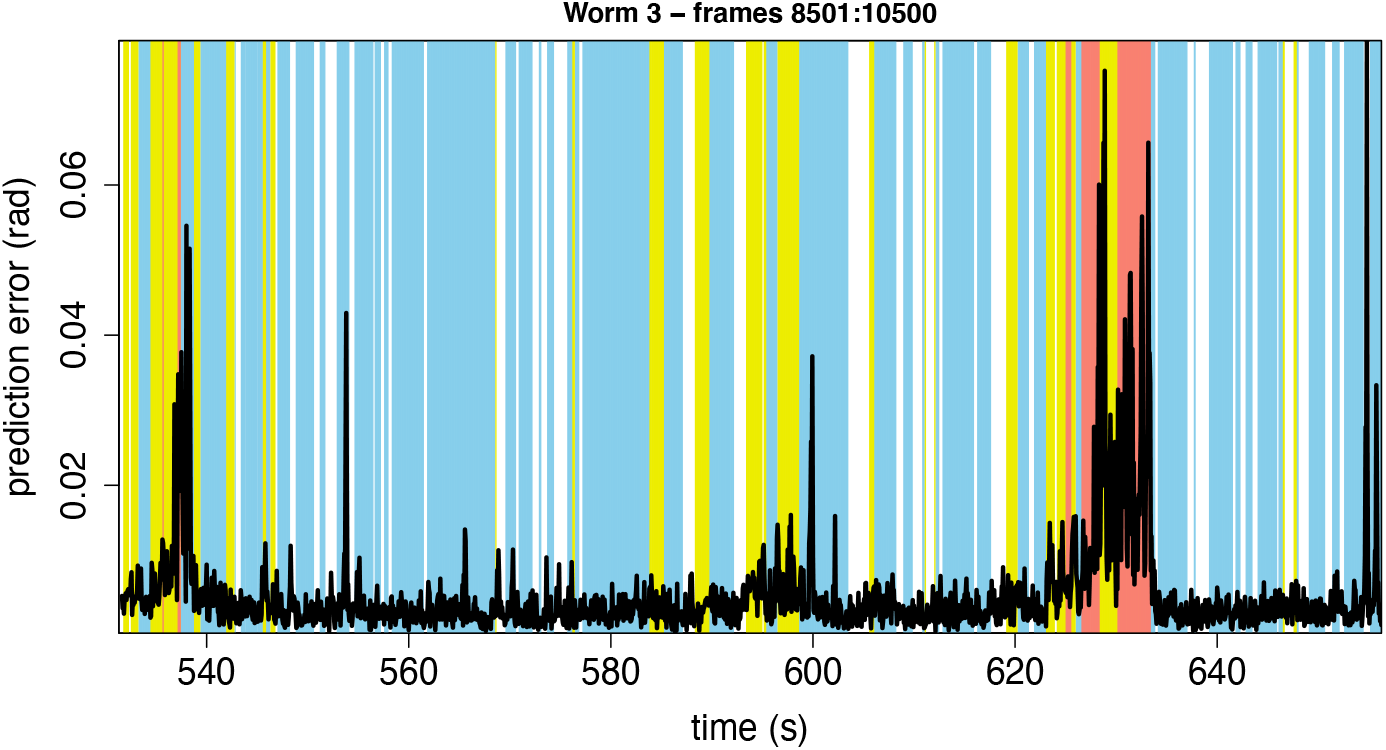
Full error time series for foraging worm 3 library (first half) and prediction (second half) intervals. Background: approximate behavioural classification (blue, forward motion; yellow, backward motion; red, tight turn).

**Appendix 2 Figure 4.**
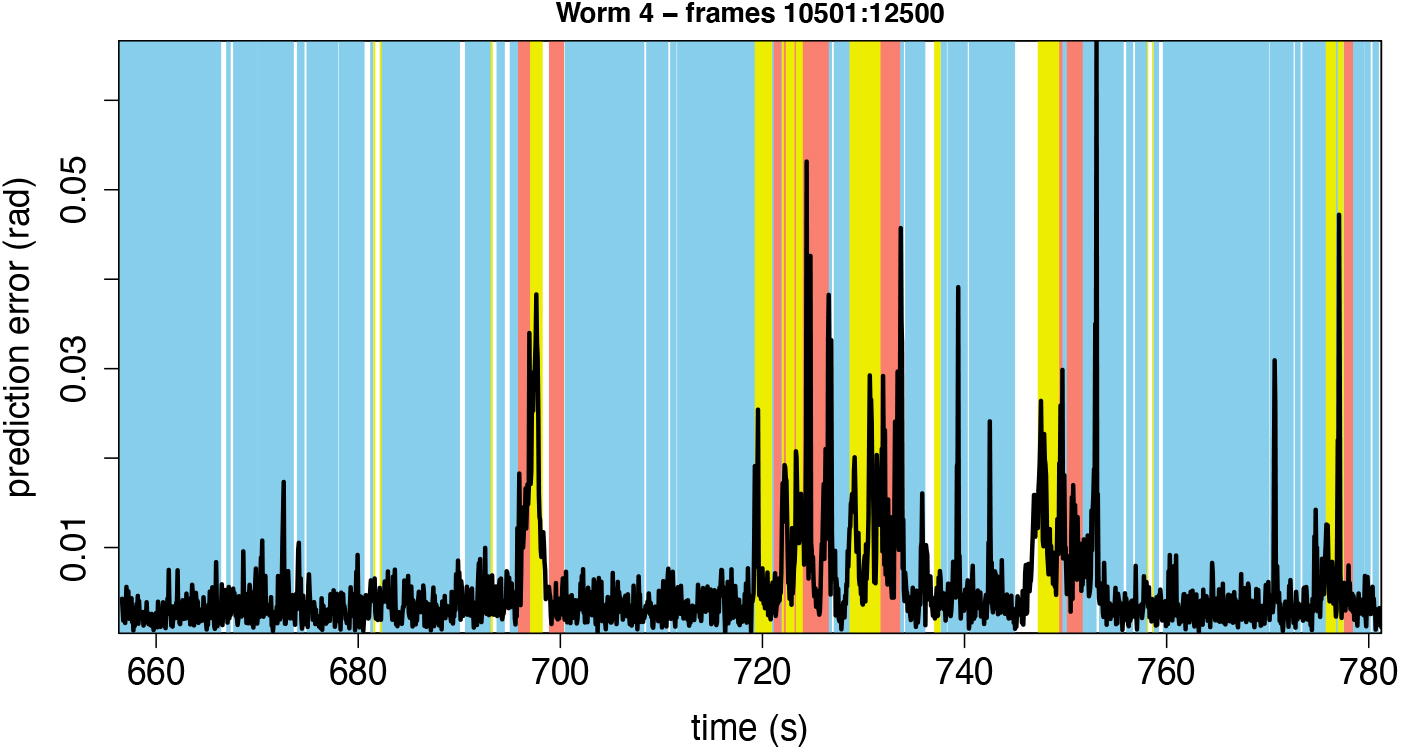
Full error time series for foraging worm 4 library (first half) and prediction (second half) intervals. Background: approximate behavioural classification (blue, forward motion; yellow, backward motion; red, tight turn).

**Appendix 2 Figure 5.**
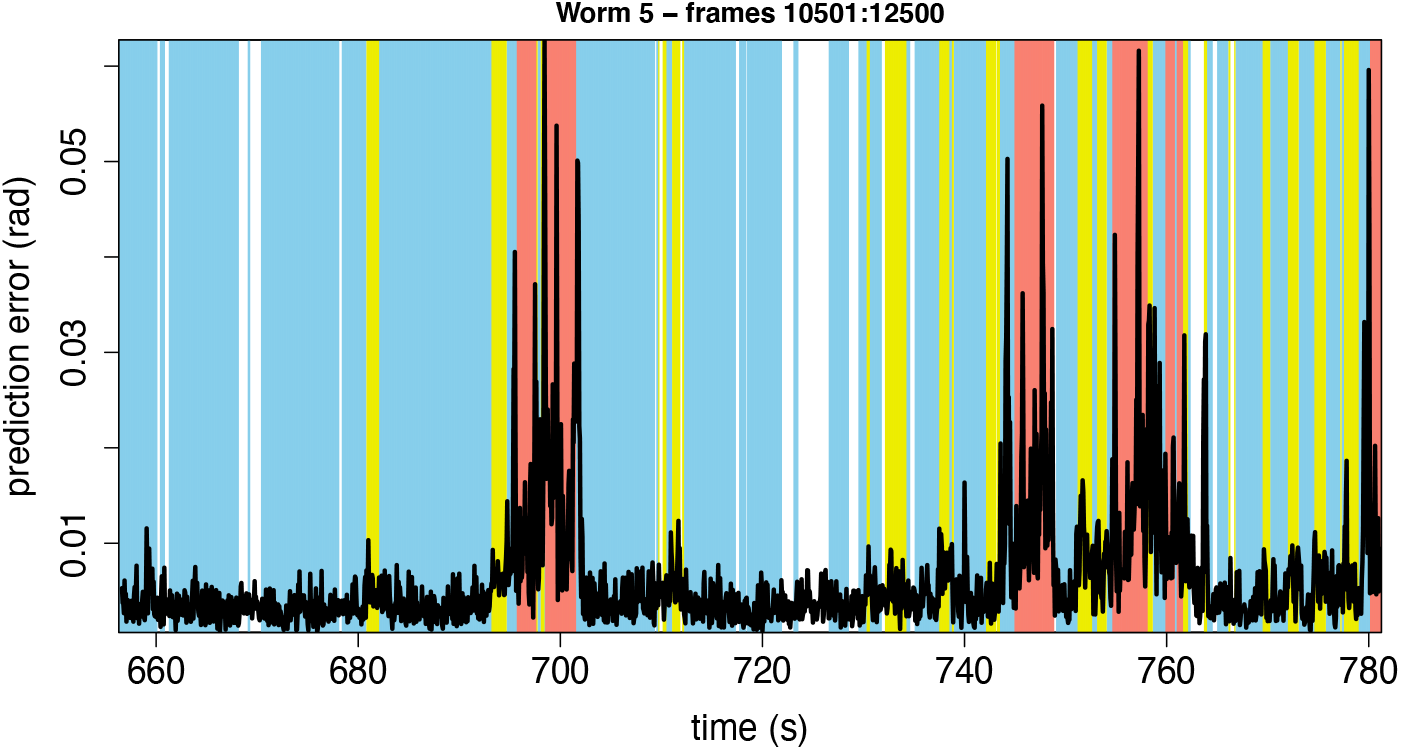
Full error time series for foraging worm 5 library (first half) and prediction (second half) intervals. Background: approximate behavioural classification (blue, forward motion; yellow, backward motion; red, tight turn).

**Appendix 2 Figure 6.**
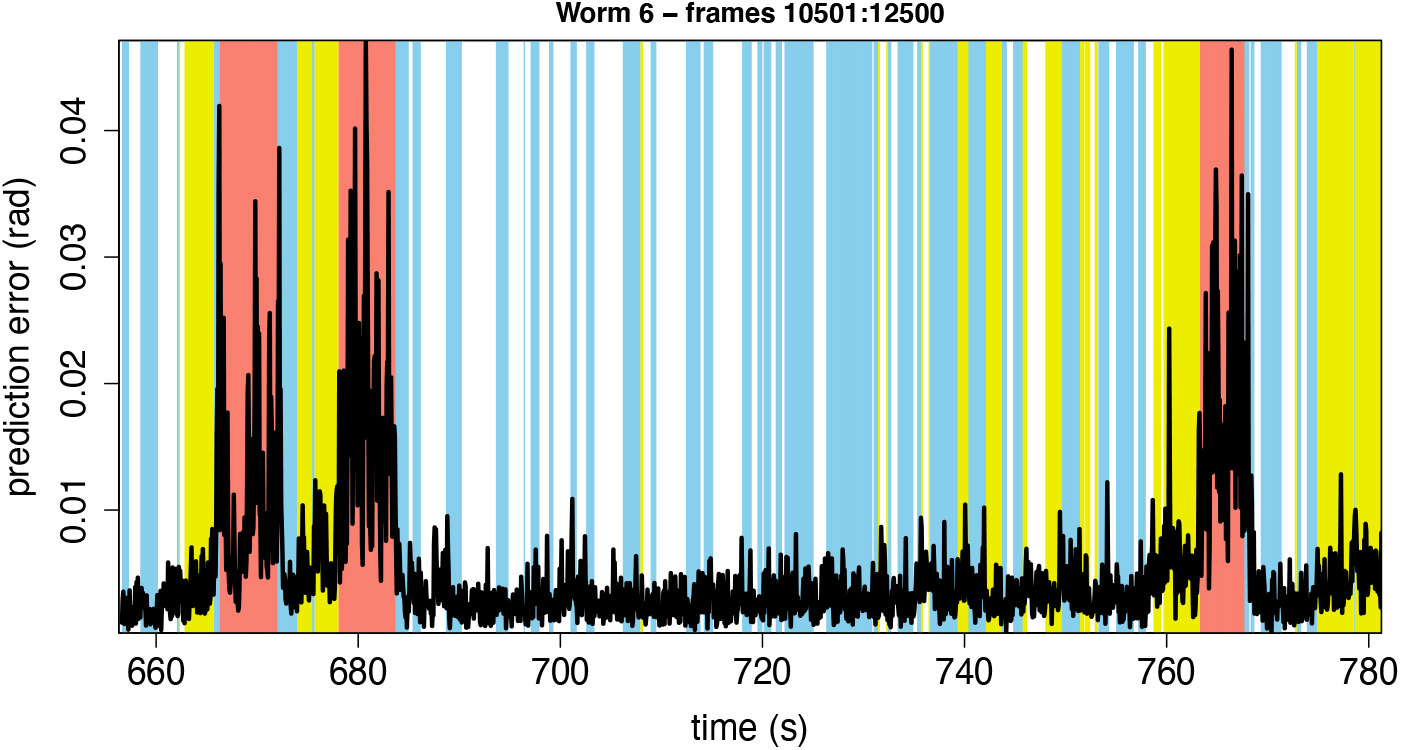
Full error time series for foraging worm 6 library (first half) and prediction (second half) intervals. Background: approximate behavioural classification (blue, forward motion; yellow, backward motion; red, tight turn).

**Appendix 2 Figure 7.**
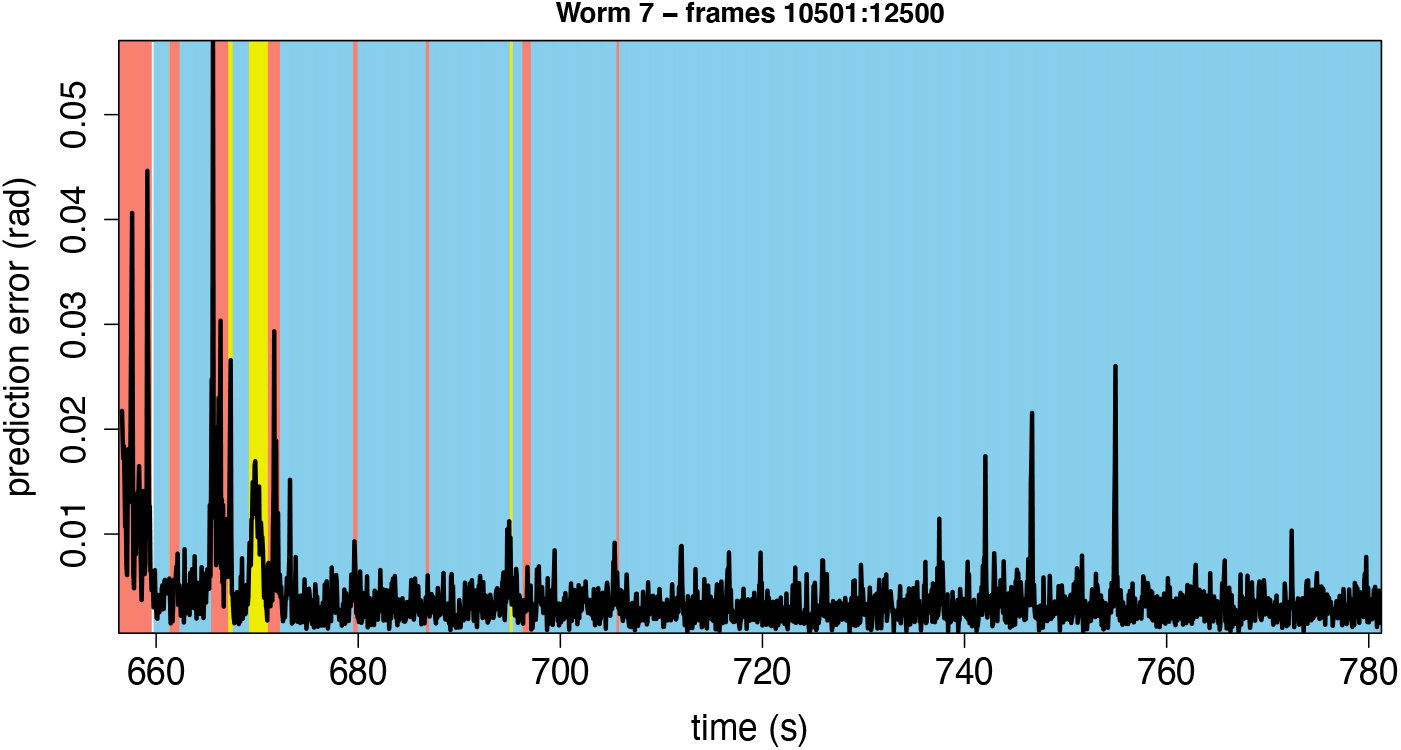
Full error time series for foraging worm 7 library (first half) and prediction (second half) intervals. Background: approximate behavioural classification (blue, forward motion; yellow, backward motion; red, tight turn).

**Appendix 2 Figure 8.**
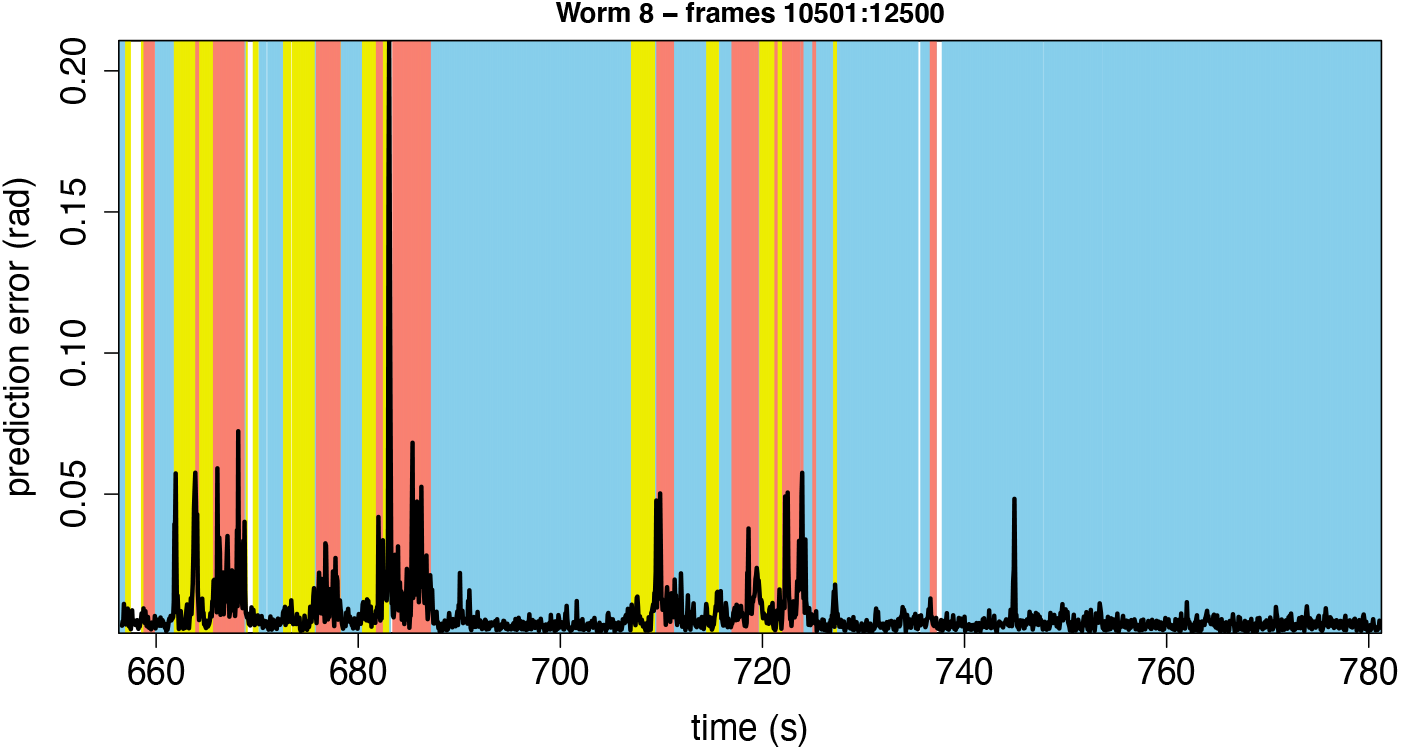
Full error time series for foraging worm 8 library (first half) and prediction (second half) intervals. Background: approximate behavioural classification (blue, forward motion; yellow, backward motion; red, tight turn).

**Appendix 2 Figure 9.**
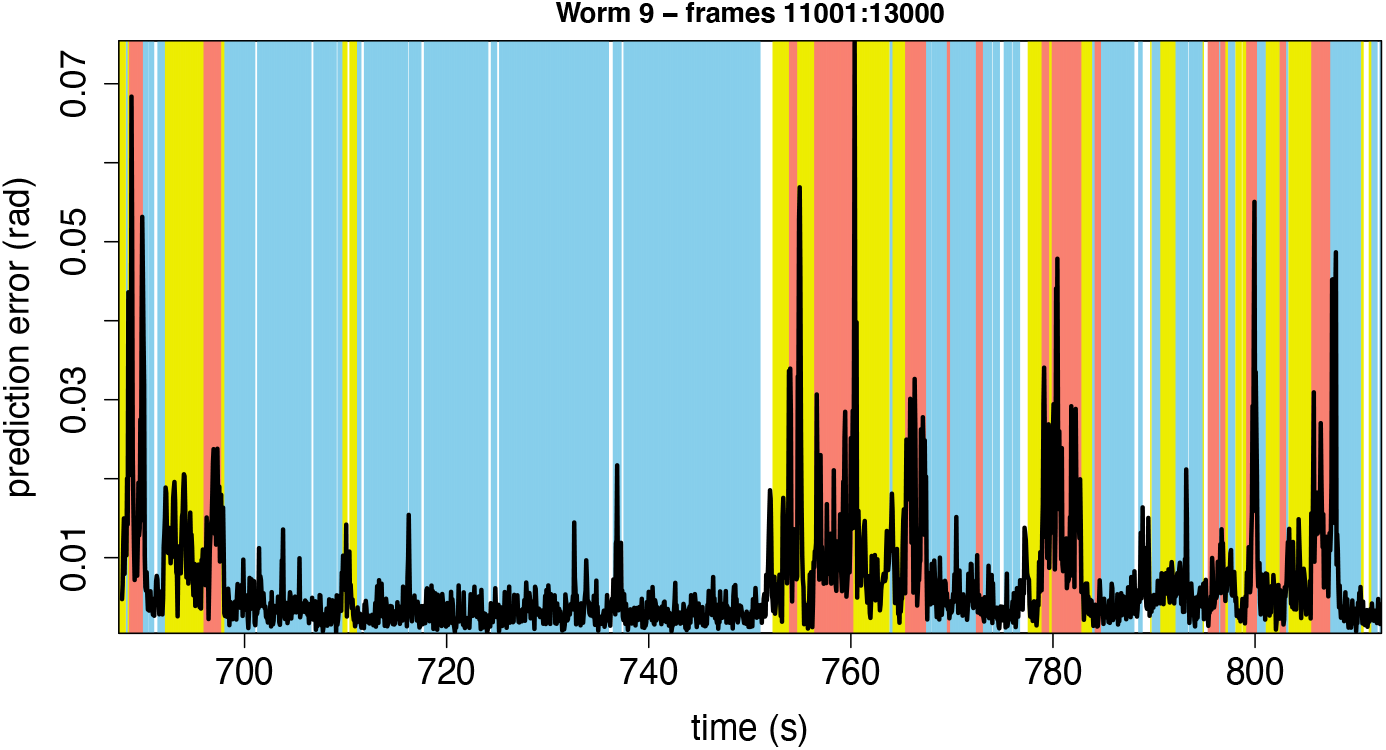
Full error time series for foraging worm 9 library (first half) and prediction (second half) intervals. Background: approximate behavioural classification (blue, forward motion; yellow, backward motion; red, tight turn).

**Appendix 2 Figure 10.**
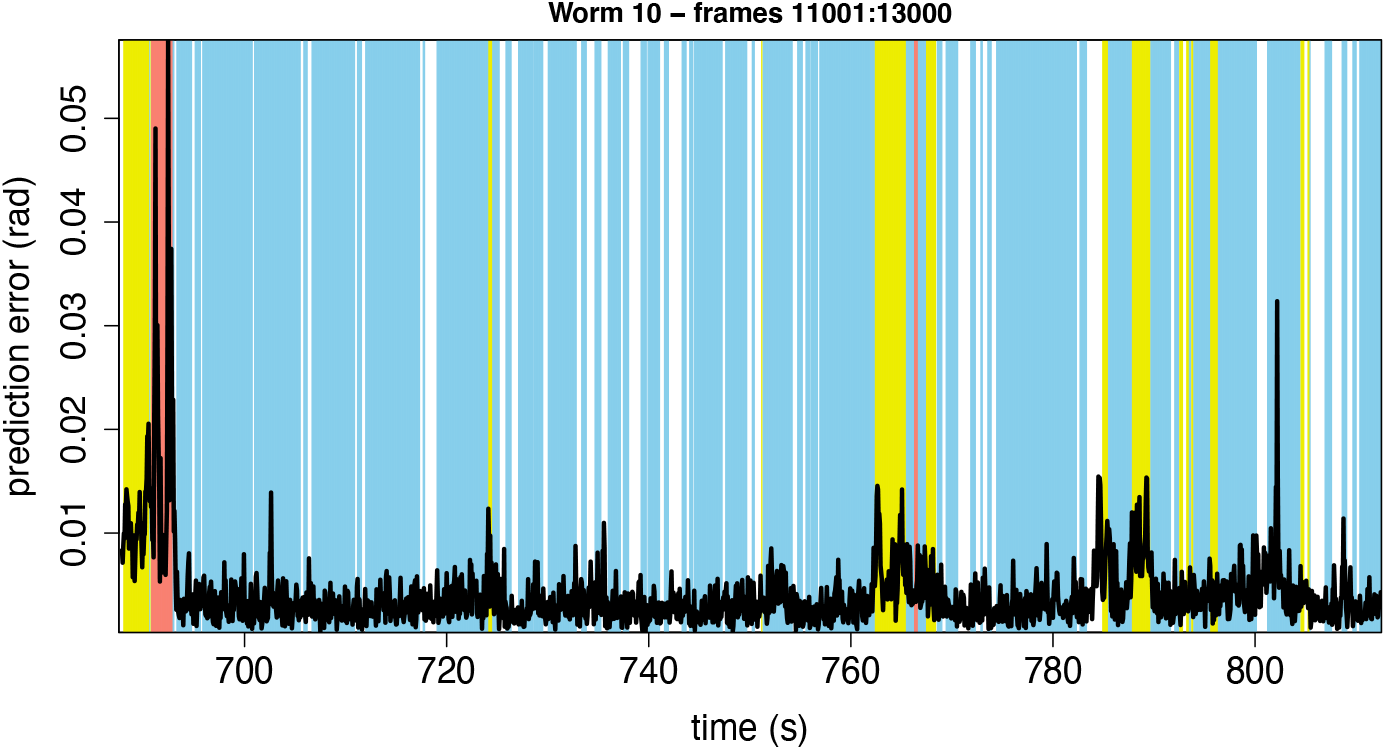
Full error time series for foraging worm 10 library (first half) and prediction (second half) intervals. Background: approximate behavioural classification (blue, forward motion; yellow, backward motion; red, tight turn).

**Appendix 2 Figure 11.**
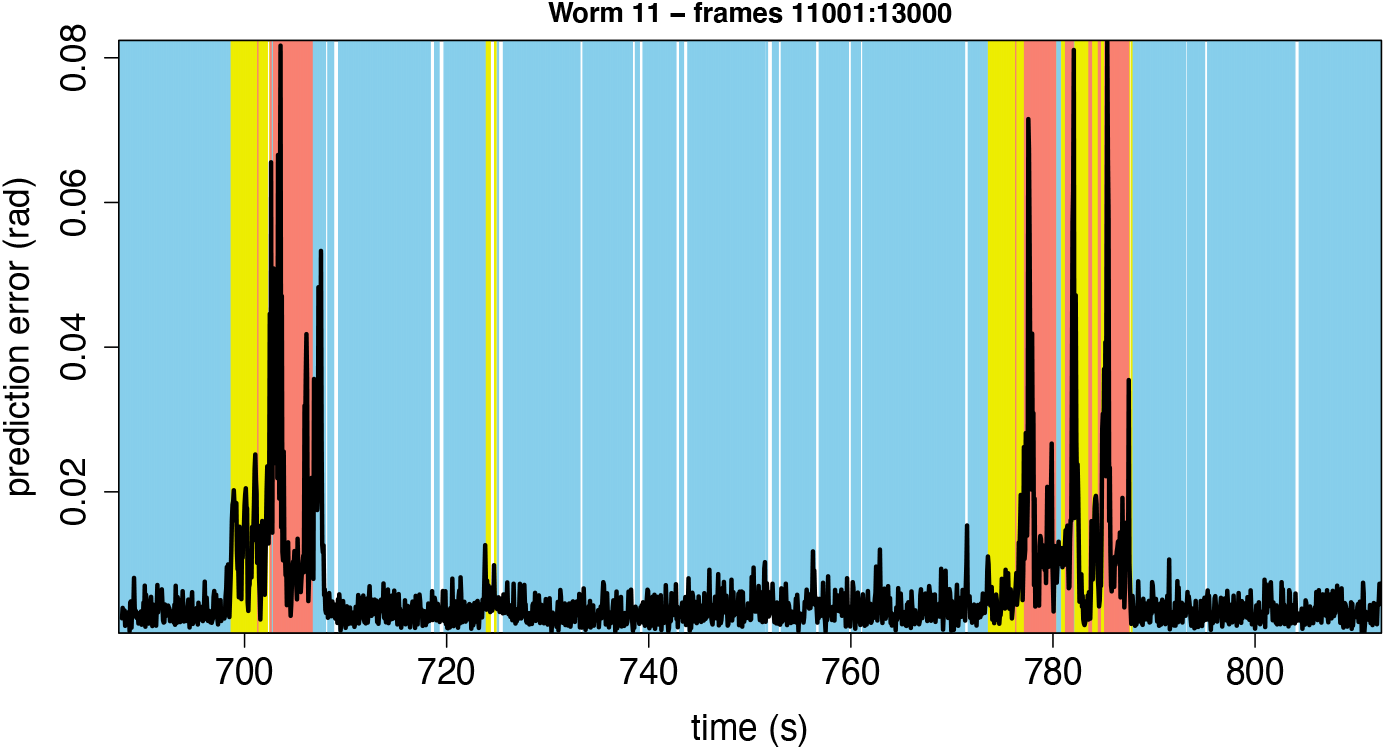
Full error time series for foraging worm 11 library (first half) and prediction (second half) intervals. Background: approximate behavioural classification (blue, forward motion; yellow, backward motion; red, tight turn).

**Appendix 2 Figure 12.**
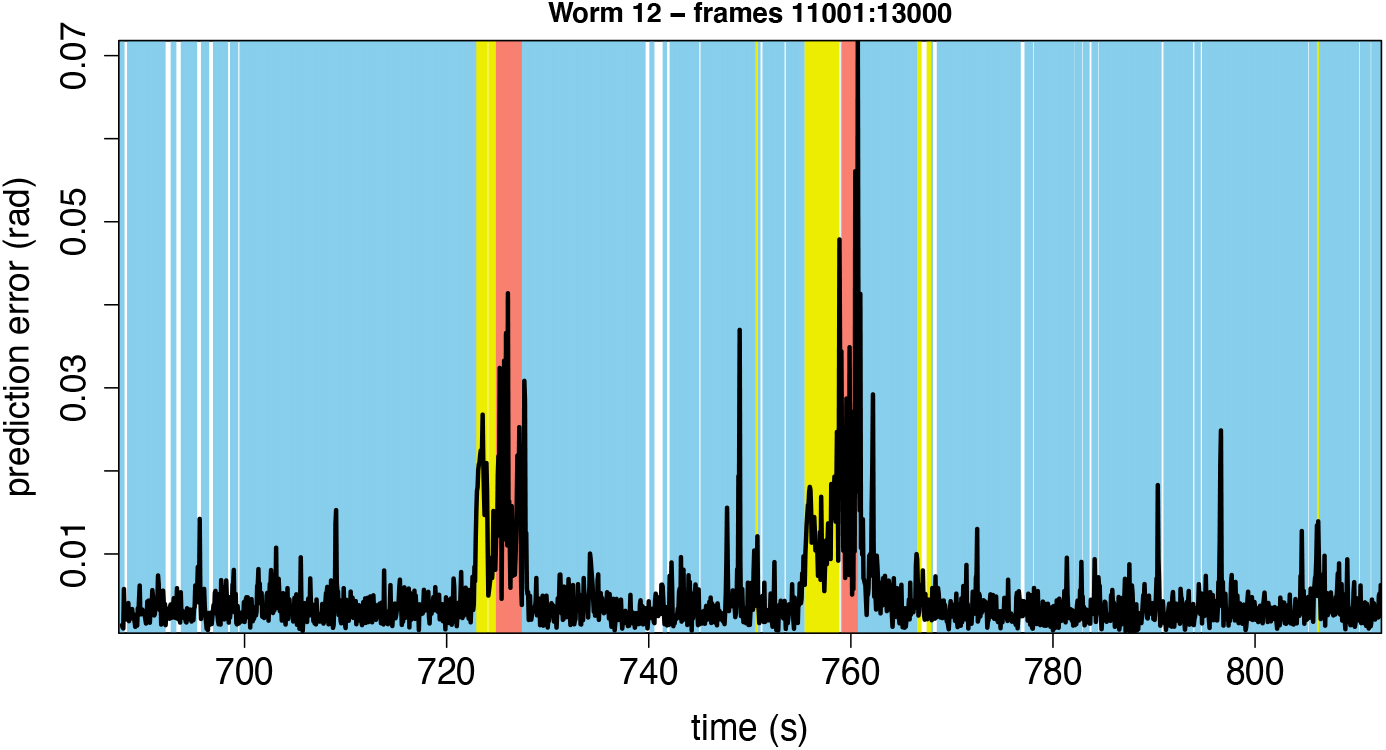
Full error time series for foraging worm 12 library (first half) and prediction (second half) intervals. Background: approximate behavioural classification (blue, forward motion; yellow, backward motion; red, tight turn).

---

**Figure 2–Figure supplement 1.**
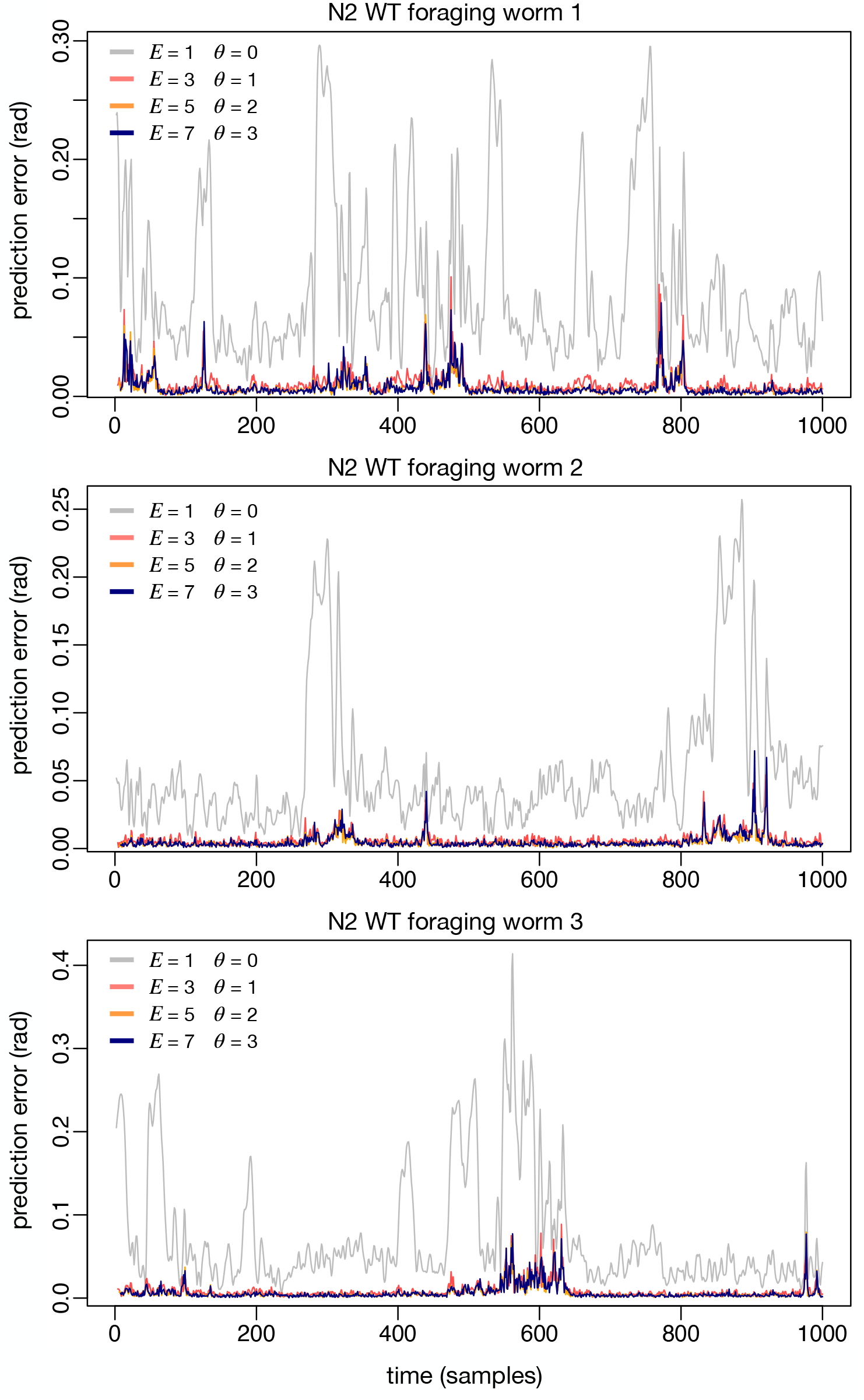
Robustness to *E* (left) and *θ* (right) for an additional wildtype sampled at 30 Hz without reconstructed self-intersecting turns. Thin lines are individual worms, thick lines are means of thin lines.

**Figure 2–Figure supplement 2.**
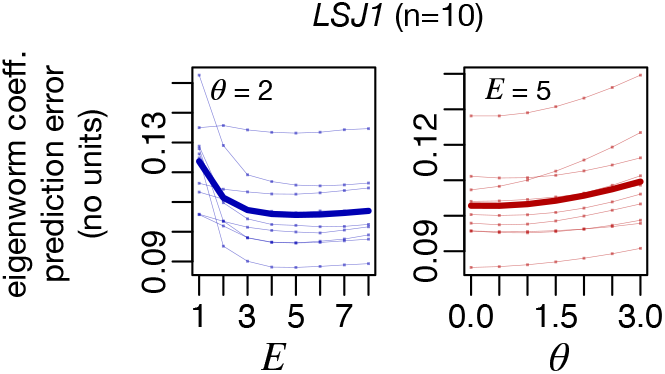
Single time series prediction error shows detailed robustness across reasonable parameter values (*E* > 1, *θ* > 0), but not for the linear case with no time delay embedding (*E* = 1, *θ* = 0). Worms correspond to first three N2 wildtype foraging worms from ***Broekmans et al. (2016)***, see Methods.

**Supplementary Table 1.**

| Mutation/strain | Recording ID |
| --- | --- |
| octr-1(ok317) (strain VC224) | tag-24 (ok371)X on food<br>R_2010_01_22__15_03_36__2__12_features |
| octr-1(ok317) (strain VC224) | tag-24 (ok371)X on food<br>L_2010_01_28__13_19_44__2__9_features |
| octr-1(ok317) (strain VC224) | tag-24 (ok371)X on food<br>L_2010_01_26__11_26__3__3_features |
| octr-1(ok317) (strain VC224) | tag-24 (ok371)X on food<br>R_2010_01_28__13_19__3__9_features |
| octr-1(ok317) (strain VC224) | tag-24 (ok371)X on food<br>L_2010_01_22__15_02_47__1__12_features |
| octr-1(ok317) (strain VC224) | tag-24 (ok371)X on food<br>R_2010_01_28__13_19_32__1__9_features |
| octr-1(ok317) (strain VC224) | tag-24 (ok371)X on food<br>R_2010_01_22__15_04_07__4__12_features |
| octr-1(ok317) (strain VC224) | tag-24 (ok371)X on food<br>R_2010_01_26__11_27_13__4__3_features |
| octr-1(ok317) (strain VC224) | tag-24 (ok371)X on food<br>R_2010_01_28__13_20_16__4__10_features |
| octr-1(ok317) (strain VC224) | tag-24 (ok371)X on food<br>L_2010_01_22__15_05_37__10_features |
| octr-1(ok317) (strain VC224) | tag-24 (ok371)X on food<br>L_2010_01_28__13_21_42__1_features |
| octr-1(ok317) (strain VC224) | tag-24 (ok371)X on food<br>R_2010_01_26__11_28_44__2_features |
| unc-80(e1069)V (strain CB1069) | unc-80 (e1069) on food<br>L_2010_04_14__14_57_39__2__12_features |
| unc-80(e1069)V (strain CB1069) | unc-80 (e1069)V on food<br>L_2009_12_09__16_36_22__2__15_features |
| unc-80(e1069)V (strain CB1069) | unc-80 (e1069)V on food<br>R_2009_12_10__13_43_33__2__8_features |
| unc-80(e1069)V (strain CB1069) | unc-80 (e1069) on food<br>L_2010_04_14__14_57__3__12_features |
| unc-80(e1069)V (strain CB1069) | unc-80 (e1069)V on food<br>R_2009_12_09__16_36__3__15_features |
| unc-80(e1069)V (strain CB1069) | unc-80 (e1069)V on food<br>R_2009_12_10__13_43__3__8_features |
| unc-80(e1069)V (strain CB1069) | unc-80 (1069)V on food<br>R_2009_12_10__13_42_50__1__9_features |
| unc-80(e1069)V (strain CB1069) | unc-80 (e1069) on food<br>L_2010_04_14__14_56_45__1__12_features |
| unc-80(e1069)V (strain CB1069) | unc-80 (e1069)V on food<br>L_2009_12_09__16_35_38__1__15_features |
| dpy-20(e1282)IV (strain CB1282) | dpy-20 (e1282)IV on food<br>L_2011_08_04__11_30_01__2__4_features |
| dpy-20(e1282)IV (strain CB1282) | dpy-20 (e1282)IV on food<br>L_2011_08_19__12_00_59__6__5_features |
| dpy-20(e1282)IV (strain CB1282) | dpy-20 (e1282)IV on food<br>R_2011_08_04__12_51_11__6__6_features |
| dpy-20(e1282)IV (strain CB1282) | dpy-20 (e1282)IV on food<br>R_2011_08_09__12_58_43__6__8_features |
| dpy-20(e1282)IV (strain CB1282) | dpy-20 (e1282)IV on food<br>R_2011_08_24__11_21_59__6__5_features |
| **dpy-20(e1282)IV (strain CB1282)** | dpy-20 (e1282)IV on food<br>R_2011_08_25__15_40_16__6__9_features |
| **dpy-20(e1282)IV (strain CB1282)** | dpy-20 (e1282)IV on food<br>L_2011_08_04__10_52_51__8__2_features |
| **dpy-20(e1282)IV (strain CB1282)** | dpy-20 (e1282)IV on food<br>L_2011_08_24__12_07_53__8__7_features |
| **dpy-20(e1282)IV (strain CB1282)** | dpy-20 (e1282)IV on food<br>L_2011_08_25__11_23_10__8__1_features |
| **dpy-20(e1282)IV (strain CB1282)** | dpy-20 (e1282)IV on food<br>R_2011_08_09__10_54_50__8__2_features |
| **dpy-20(e1282)IV (strain CB1282)** | dpy-20 (e1282)IV on food<br>R_2011_08_19__12_43_48__8__7_features |
| **dpy-20(e1282)IV (strain CB1282)** | dpy-20 (e1282)IV on food<br>L_2011_08_09__10_33_28__7__1_features |
| **Wildtype strain LSJ1** | 300 LS51 on food<br>L_2010_11_26__16_25__3__13_features |
| **Wildtype strain LSJ1** | 300 LS51 on food<br>L_2011_02_17__11_36__3__3_features |
| **Wildtype strain LSJ1** | 300 LS51 on food<br>R_2010_11_26__11_47__3__5_features |
| **Wildtype strain LSJ1** | 300 LS51 on food<br>L_2011_02_17__12_48__3__7_features |
| **Wildtype strain LSJ1** | 300 LS51 on food<br>L_2010_11_26__10_38_21__7__1_features |
| **Wildtype strain LSJ1** | 300 LS51 on food<br>R_2011_02_25__10_31_11__7__1_features |
| **Wildtype strain LSJ1** | 300 LSJ1 on food<br>L_2011_03_10__10_52_16__6__1_features |
| **Wildtype strain LSJ1** | 300 LSJ1 on food<br>R_2011_03_07__11_42_35__6__3_features |
| **Wildtype strain LSJ1** | 300 LSJ1 on food<br>R_2011_04_12__12_30_20__6__5_features |
| **Wildtype strain LSJ1** | 300 LSJ1on food<br>R_2011_03_15__12_37_50__6__5_features |

## Notes

### Competing Interest Statement

The authors have declared no competing interest.

